# Elastic Network Modeling of Cellular Networks Unveils Sensor and Effector Genes that Control Information Flow

**DOI:** 10.1101/2021.06.04.446730

**Authors:** Omer Acar, She Zhang, Ivet Bahar, Anne-Ruxandra Carvunis

## Abstract

The high-level organization of the cell is embedded in long-range interactions that connect distinct cellular processes. Existing approaches for detecting long-range interactions consist of propagating information from source nodes through cellular networks, but the selection of source nodes is inherently biased by prior knowledge. Here, we sought to derive an unbiased view of long-range interactions by adapting a perturbation-response scanning strategy initially developed for identifying allosteric interactions within proteins. We deployed this strategy onto an elastic network model of the yeast genetic network. The genetic network revealed a superior propensity for long-range interactions relative to simulated networks with similar topology. Long-range interactions were detected systematically throughout the network and found to be enriched in specific biological processes. Furthermore, perturbation-response scanning identified the major sources and receivers of information in the network, named effector and sensor genes, respectively. Effectors formed dense clusters centrally integrated into the network, whereas sensors formed loosely connected antenna-shaped clusters. Long-range interactions between effector and sensor clusters represent the major paths of information in the network. Our results demonstrate that elastic network modeling of cellular networks constitutes a promising strategy to probe the high-level organization of the cell.

## Introduction

Cellular networks are high-level representations of the relationships between genes or between their encoded products. These networks represent genes as nodes and interactions as edges. The interactions may involve direct physical relationships between biomolecules (proteins, transcripts), or functional relationships between genes including epistatic genetic interactions or coordinated regulation of gene expression.^1^ In-depth analysis of the local interactions around one or several genes of interest allows to identify biological modules^2^ and disease-associated groups of genes,^3^ and to elucidate unknown gene functions.^4,5^ Taken in concert, global analysis of the structure and dynamics of cellular networks can aid in further understanding the overarching biological and physical mechanisms that govern cellular machinery and behavior.^6^

Genetic interactions play a central role in genotype-phenotype relationships.^7^ Genetic perturbations (e.g., gene deletions or mutations) may alter only the local interaction neighborhood for a molecule, but the effects of a local alteration can also propagate through the network and cause changes on a larger scale. For instance, genetic alterations that rewire an established transcriptional program, disrupt chromatin context, or prevent the activation of a signal transduction pathway can impact numerous downstream genes and processes.^8^ Therefore, to capture the breadth of genetic perturbation effects, it is crucial to study both short- and long-range interactions.^9^

The study of long-range interactions poses a computational challenge.^10^ Network propagation (also referred to as information transfer or geometric diffusion) methods^11–13^ have been widely used to identify long-range relationships between genes^14–16^ or within biomolecular structures.^17^ The basic principle of these methods is to model a diffusion process starting from a source node, similar to the flow of a liquid or heat in a solid matter, and to calculate the amount of diffusion often modeled as a Markovian process across the network. The amount of diffusion across the network is used as a metric quantifying the long-range relationship. For some applications, this is equivalent to a random walk with restart process on the network nodes.^16^ These propagation methods have a wide range of applications from identifying disease-related genes^18,19^ to protein homology detection.^20^

An important caveat for the use of network propagation for genetic networks is the requirement for prior information about well-characterized source genes, such as disease genes. This introduces an inherent bias that prevents the discovery of novel relationships that are not related to prior knowledge. Thus, to obtain a comprehensive understanding of a network’s long-range relationships, an unbiased approach is needed in which all nodes should be considered as possible sources and all possible long-range relationships should be investigated. However, not all genes will engage equally in long-range relationships. Based on their biological properties, some genes could be involved in many cellular pathways and thus might be more effective at propagating information to other genes; or some genes might be involved in specific signaling pathways such that their sensitivity at receiving and integrating signals is crucial to their cellular role. Thus, unbiased identification of the key propagation-mediating genes is critical to discover important long-range relationships in genetic networks.

To achieve this goal, we leveraged a perturbation-response scanning (PRS) strategy initially developed for the unbiased identification of long-range interactions within molecular structures.^21–23^ The structures (proteins or chromosomes) are represented by elastic network models (ENM)^24^ where each network node represents a physical entity (e.g., a residue, domain, monomer, or gene locus in the chromatin)^25^ and each network edge is modeled as a spring that represents a physical interaction between the nodes. ENM representation allows for the application of forces/perturbations on network nodes and then measurement of the cooperative motions/responses of all other nodes, where the former represent the initial information and the latter represents the propagated information (Figure 1A). We reasoned that PRS could be successfully extended to genetic networks because they lend themselves to ENM representations and because spring-based modeling of genetic networks has already proven to be valuable for both visualization^26^ and biological inference.^27^

**Figure 1:**
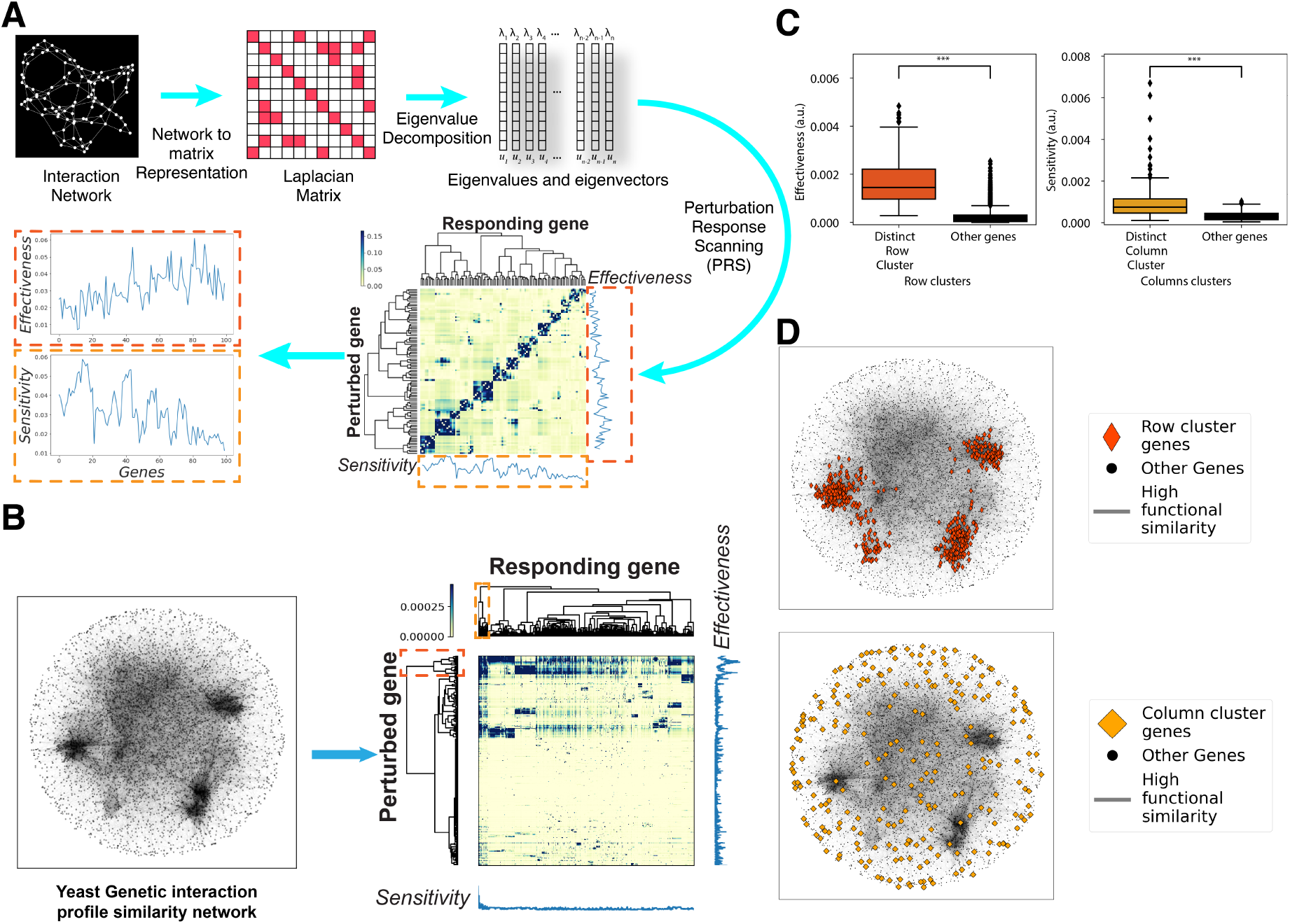
Perturbation Response Scanning (PRS) in the yeast genetic interaction profile similarity network (GI PSN). A) The PRS strategy. A network is first transformed into the Laplacian matrix describing the network connectivity. Eigenvalue decomposition of the Laplacian yields the eigenvalues and eigenvectors used to calculate the PRS matrix. Each row of the PRS matrix corresponds to the perturbed node, each column corresponds to responding nodes and the colors show the magnitude of the response. Row and column averages of the PRS matrix represent the effectiveness (*right ordinate*) and sensitivity (*lower abscissa*) profiles, respectively. B) PRS analysis of the GI PSN (*left*) yields the PRS matrix shown on the *right*. The nodes on the network are the genes and the edges represent high profile similarity. This network representation was used throughout the paper. *Dashed boxes* on the dendrograms along two axes of the PRS matrix indicate distinct row and column clusters. C) Effectiveness (*left*) and sensitivity (*right*) boxplots showing the differences between row and column clusters, respectively (***: *p* < .001, a.u.: arbitrary units, also used for the remaining figures). D) Representation of the distinct row (*top*) and column (*bottom*) clusters within the GI PSN.

Here, we adapted the PRS strategy to identify critical propagation-mediating nodes and obtain a global, unbiased view of long-range interactions in the comprehensive genetic interaction profile similarity network (GI PSN) generated for *S. cerevisiae* by Costanzo *et al*.^28^ We evaluated the signal propagation ability of each yeast gene using two metrics: sensitivity and effectiveness. Sensitivity is defined as the propensity to receive information, independent of the source; effectiveness is defined as the ability to transmit information to other genes.^22^ Genes distinguished by their high ability to receive and transmit information are defined as sensors and effectors, respectively. Our analysis uncovers critical network clusters formed by effector and sensor genes and unveils the long-range interactions connecting seemingly unrelated cellular processes.

## Results

### PRS clusters genes based on their potential to receive and transmit information

The GI PSN contains 5,183 genes and 39,816 edges representing functional similarity between genes (Figure 1B, *left*). We constructed an ENM representation of the GI PSN and applied the PRS strategy to this network by perturbing each gene individually and measuring the responses of the other genes. This resulted in a 5,183-by-5,183 PRS matrix (Figure 1B, *right*) representing the perturbation-response relationship between all pairs of genes. Hierarchical clustering of the PRS matrix rows and columns clearly delineated groups of genes based on their information propagation profiles. Notably, one row and one column cluster were separated from the rest of the genes in the dendrograms (Figure 1B, *dashed boxes* on the dendrograms). The nodes in these two distinct clusters displayed higher effectiveness and sensitivity than the rest of the genes in the network, respectively (Figure 1C, *p* <.01, permutation test). Next, we mapped the genes belonging to these distinct clusters on the network. The distinct row cluster corresponded to highly connected, central regions of the network. In contrast, the distinct column cluster corresponded to genes that are distributed throughout the network, with a tendency to be in peripheral locations (Figure 1D). Overall, PRS-based clustering identified two classes of genes; one with high effectiveness located in densely connected regions; and another with high sensitivity, at loosely connected regions of the network.

### GI PSN displays a remarkable potential for long-range interactions

The local connectivity of each node can be summarized by its degree (number of neighbors), and the behavior of a network is largely characterized by its degree distribution.^29^ Thus, we sought to understand how the degree of nodes and the degree distribution of the GI PSN influence effectiveness and sensitivity profiles. We found that effectiveness was highly correlated with degree (Figure 2A, *R* = .9), whereas sensitivity was not (*ρ* = −.028), although nodes with low degrees (degree<10) tended to show higher sensitivity (Figure 2B, *ρ* = −.97 for degree<10). To investigate the significance of these results, we compared the GI PSN to random networks generated by rewiring the GI PSN edges while keeping the degree distribution constant (Figure 2C). We first compared correlations derived from the GI PSN to those derived from 100 randomly rewired networks. The results showed that the GI PSN had a significantly stronger degree effectiveness correlation and weaker degree-sensitivity correlation than rewired networks (Figure 2D-E, *p* < .001, empirical *p*-value). Next, we examined the network’s effectiveness and sensitivity distributions of the real and rewired networks. Nodes in the GI PSN had overall higher effectiveness and sensitivity values than the random networks (Figure 2F-G, *p* < .001, empirical *p*-value). However, the shapes of both the distributions of effectiveness and sensitivity bear some interesting resemblance between the real and the randomized networks (Figure 2F-G, *red dotted and cyan solid curves*). Since effectiveness and sensitivity measure the potential of the nodes to transmit and receive information, respectively, our results demonstrate that the GI PSN harbors significantly stronger signal propagation propensities than expected from its degree distribution alone.

**Figure 2:**
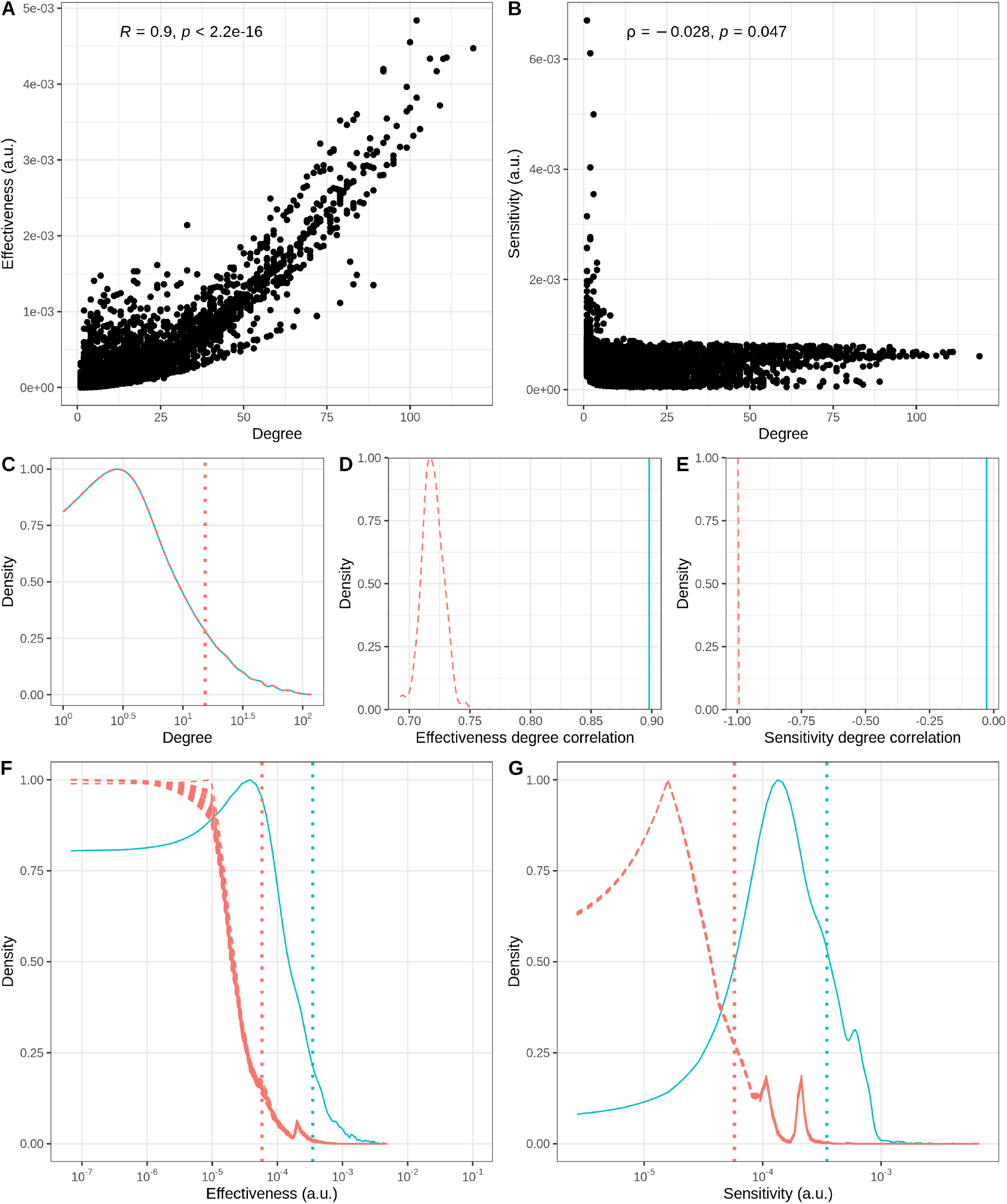
The GI PSN displays superior information propagation potential compared to randomly rewired networks with identical degree distributions. A) Degree and effectiveness scatter plot shows strong correlation between degree and effectiveness in the GI PSN (Pearson correlation, *R* = .9). B) Degree and sensitivity scatter plot shows no correlation between degree and sensitivity is observed in the GI PSN (Spearman’s rank correlation, *ρ* = −.028). C) Degree distributions for the GI PSN (*cyan*) and the rewired networks (*red*). These distributions overlap by design. D) The correlation between degree and effectiveness is significantly higher in the GI PSN (*blue vertical line*) than that expected for the rewired networks (*dashed red distribution*, average R=.72). E) The correlation between degree and sensitivity is significantly weaker in the GI PSN (*blue vertical line*) than expected from rewired networks (*dashed red distribution*, average *ρ* = −.99). Nodes in the GI PSN (*blue distributions*) exhibit significantly higher effectiveness (F) and sensitivity (G) compared to random network nodes (*red dashed distributions*).

### Sensors form “antenna-shaped” biological clusters loosely connected with the GI PSN

Genes with higher sensitivity are more likely to be involved in long-range interactions due to their ability to integrate information from other parts of the network. Thus, we first defined genes with high sensitivity (top 1%) as sensor genes (n=52) and investigated their topological and biological properties. Sensors tended to have low degrees and, in many cases, had only a single connection (Figure 2B). We hypothesized that sensors may be directly connected to genes with high effectiveness, as was observed for protein structure networks.^22^ However, this was not the case in the GI PSN. Sensors tended to be connected to other low degree genes (Figure 3A) while the genes with high effectiveness all had high degrees (Figure 2A). In fact, the first neighbors of sensors had degrees about two orders of magnitude smaller than the first neighbors of non-sensor genes (Figure 3A). Next, we investigated whether the sensors are connected to each other more than expected given their low degree. We compared the sensors to randomly sampled nodes with the same degree and calculated the percentage of the number of the between-group edges to the total number of edges. We found that the sensors had a strong tendency to connect to each other (Figure 3B, ∼224 fold, *p* < .001, empirical *p*-value), revealing the existence of sensor clusters.

**Figure 3:**
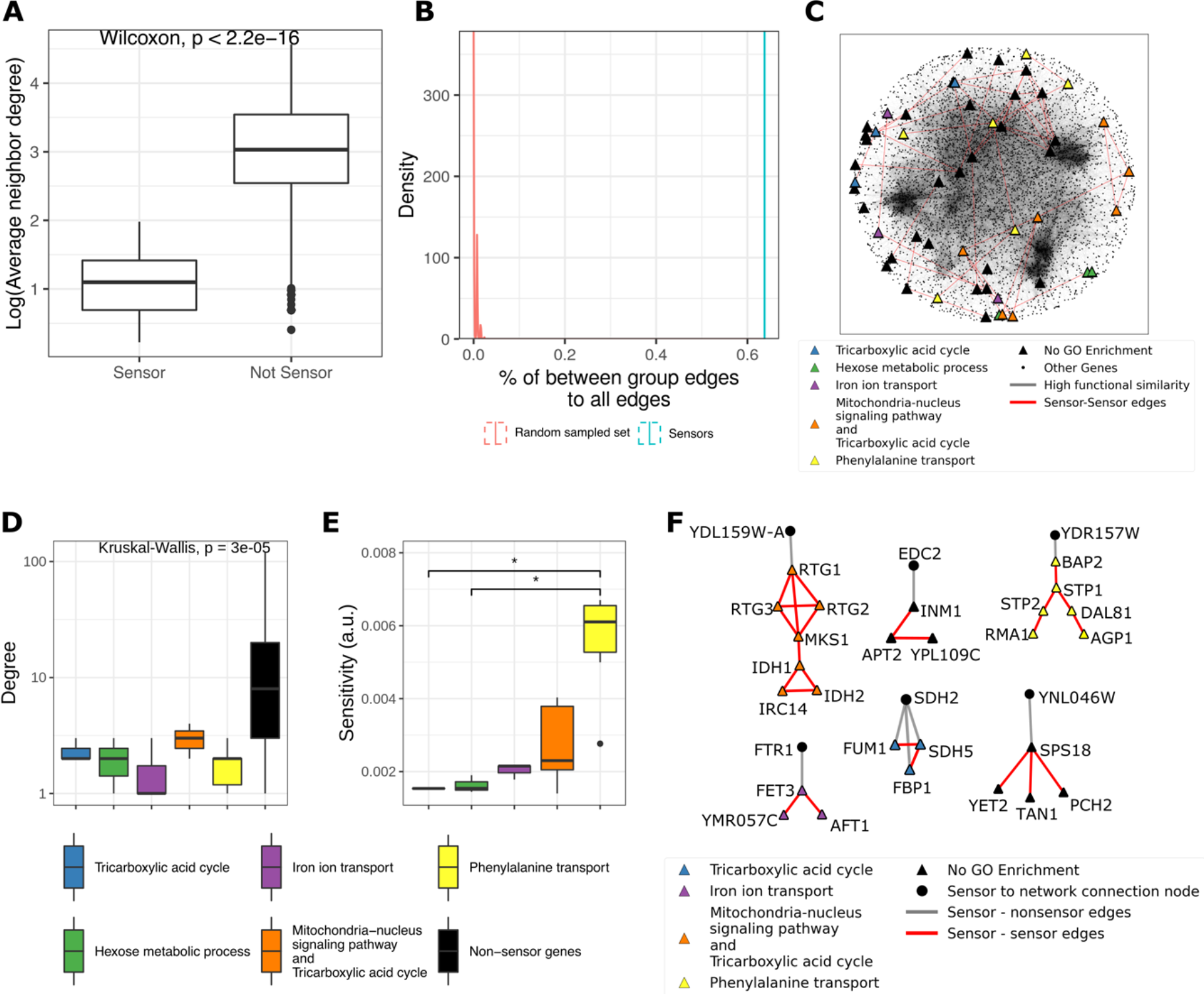
Sensors form biologically enriched low-degree gene clusters on the network periphery. A) The first neighbors of sensors have low average degree relative to the neighbors of other genes in the network. B) Sensors are more densely connected to each other than expected from randomly sampled nodes with the same degree, as measured by the percentage of the between-group edges to total edges the nodes have (sensor group: *blue vertical line*; groups of randomly sampled nodes with same degree: *red distribution*). C) Representation of sensor genes in the GI PSN. Node colors represent sensor clusters with distinct GO term enrichments (node colors and shapes, and edge colors were used similarly for the following figures). D) Sensor clusters exhibit comparable degrees (*colored left*) and are significantly lower than the degrees of other genes in the network (Kruskal-Wallis, group-wise comparison including non-sensor genes). E) The sensor groups have similar sensitivity, except for ‘phenylalanine transport’ related sensors which show higher sensitivity (*: *p* < .05, for corrected *p*-values calculated by Mann-Whitney). F) Sensor clusters showing antenna motifs. Each cluster is shown with the network node that connects the cluster to the rest of the network (*Triangles*: Sensors, *Circles*: non-sensor connecting node)

The sensors could be separated into nine clusters composed of connected components of three or more sensor genes which include 41 sensors out of the 52 total sensors in the network. Five of these nine sensor clusters could be assigned to a specific biological process through gene ontology (GO) enrichment analysis: tricarboxylic acid cycle (TCA) cycle, hexose metabolic process, iron ion transport, mitochondria-nucleus signaling, and phenylalanine transport (Figure 3C, **Supplementary Data 1**). While these sensor clusters had a lower degree than other genes in the network (Figure 3D, *p* < .001, Kruskal-Wallis, group-wise comparison including non-sensor genes), we did not see significant differences in the average node degree between these clusters (Figure 3D, *p* = .16, Kruskal-Wallis, group-wise comparison excluding non-sensor genes). Intriguingly, the sensor cluster related to ‘phenylalanine transport’ displayed the highest sensitivity (Figure 3E).

To understand what distinguishes sensors from the many other low-degree genes in the network that did not display high sensitivity, we studied their topologies in depth. Interestingly, we found that most sensors were connected to the rest of the network by a single non-sensor node, creating antenna-shaped motifs (Figure 3F). These antenna motifs appeared to form an information bottleneck where the perturbation signal can enter the sensor cluster but cannot escape easily and transfer the signal to other nodes outside of the cluster. Thus, the sensitivity of lower degree nodes within antenna motifs, as opposed to those outside the motifs, may be increased by the local accumulation of PRS signals.

### Effectors form biological clusters centrally integrated within the GI PSN

To investigate the most influential genes in the network, we defined genes with high effectiveness (top 1%) as effector genes (n=52) and studied their topological and biological properties. As opposed to sensors, effectors tended to connect to other high degree genes (Figure 4A). However, effector-effector edges consisted of only 7% of all edges involving effectors due to their extremely high degree. Nevertheless, they formed distinct network clusters, with no direct connections between different effector clusters. We could separate all 52 effectors into three connected components. Each effector cluster could be assigned to a specific biological process by GO enrichment analysis: respiratory complex assembly, Golgi vesicle transport, and chromosome segregation (Figure 4B, **Supplementary Data 1**). We found that all three clusters have significantly higher average degrees than other genes in the network (Figure 4C, *p* < .001, Kruskal Wallis, group-wise comparison including non effector genes) and effectors involved in Golgi vesicle transport have a slightly but significantly higher average degree than effectors from the other two effector clusters (Figure 4C, *p* < .001, Kruskal Wallis, group-wise comparison excluding non-effector genes). However, there was no significant difference in effectiveness values between the three clusters (Figure 4D, *p* = 0.36, Kruskal Wallis). In summary, effectors formed three biological clusters that are centrally integrated within the GI PSN while being clearly distinct from each other.

**Figure 4:**
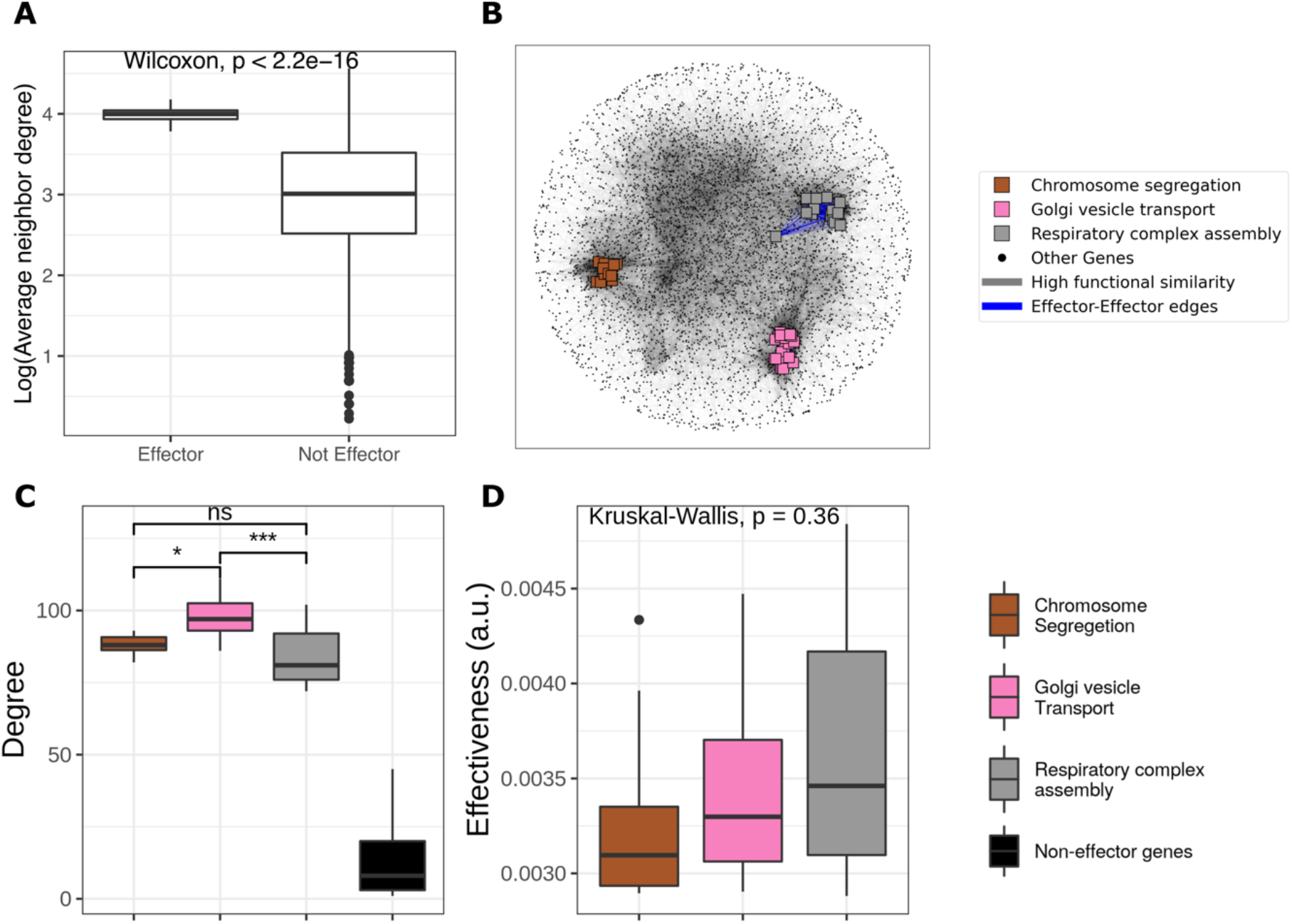
Effectors form biologically enriched high degree gene clusters at the center of the network. A) The neighbors of effectors have high average degree relative to the neighbors of other genes in the network. B) Representation of effector genes in the GI PSN. Node colors represent effector clusters with distinct GO term enrichments (node colors and shapes, and edge colors were used similarly for the following figures). C) Effectors have a higher degree than other genes and the effector cluster with ‘Golgi vesicle transport’ enrichment has a higher average degree than other effector clusters (ns: non-significant, *: *p* < .05, ***: *p* < .001, for corrected *p*-values calculated by Mann-Whitney). D) Effectiveness values are not significantly different for different clusters of effectors.

### Systematic detection of long-range interactions in the GI PSN

The PRS matrix (Figure 1B) quantifies information propagation between all pairs of genes in the GI PSN. The strongest long-range interactions can be extracted systematically by identifying the genes with the strongest response to each perturbed gene, and the genes causing the strongest perturbation to each responding gene. To evaluate the biological relevance of these systematically extracted long-range interactions, we created distinct sets of ranked ‘highly responsive’ and ‘highly influential’ genes based on their PRS profiles and evaluated the functional relatedness of genes within each set. First, for each row of the PRS matrix, or each gene acting as the perturbing source, we defined the genes that showed the highest responses on that row as the set of ‘highly responsive’ genes specific to the perturbed gene (Figure 5A, *top*). This method of ranking genes based on their responsiveness to a perturbed source gene has been used to identify disease-related genes.^19,30,31^ Additionally, we also implemented a novel target-based ranking procedure. For each column of the PRS matrix, or each gene acting as the responding target, we defined the genes that induced the highest perturbations on that column as the set of ‘highly influential’ genes specific to the responding gene (Figure 5A, *bottom*). We then performed GO term enrichment analysis for ‘highly responsive’ or ‘highly influential’ sets which were defined separately for each source and target gene.

**Figure 5:**
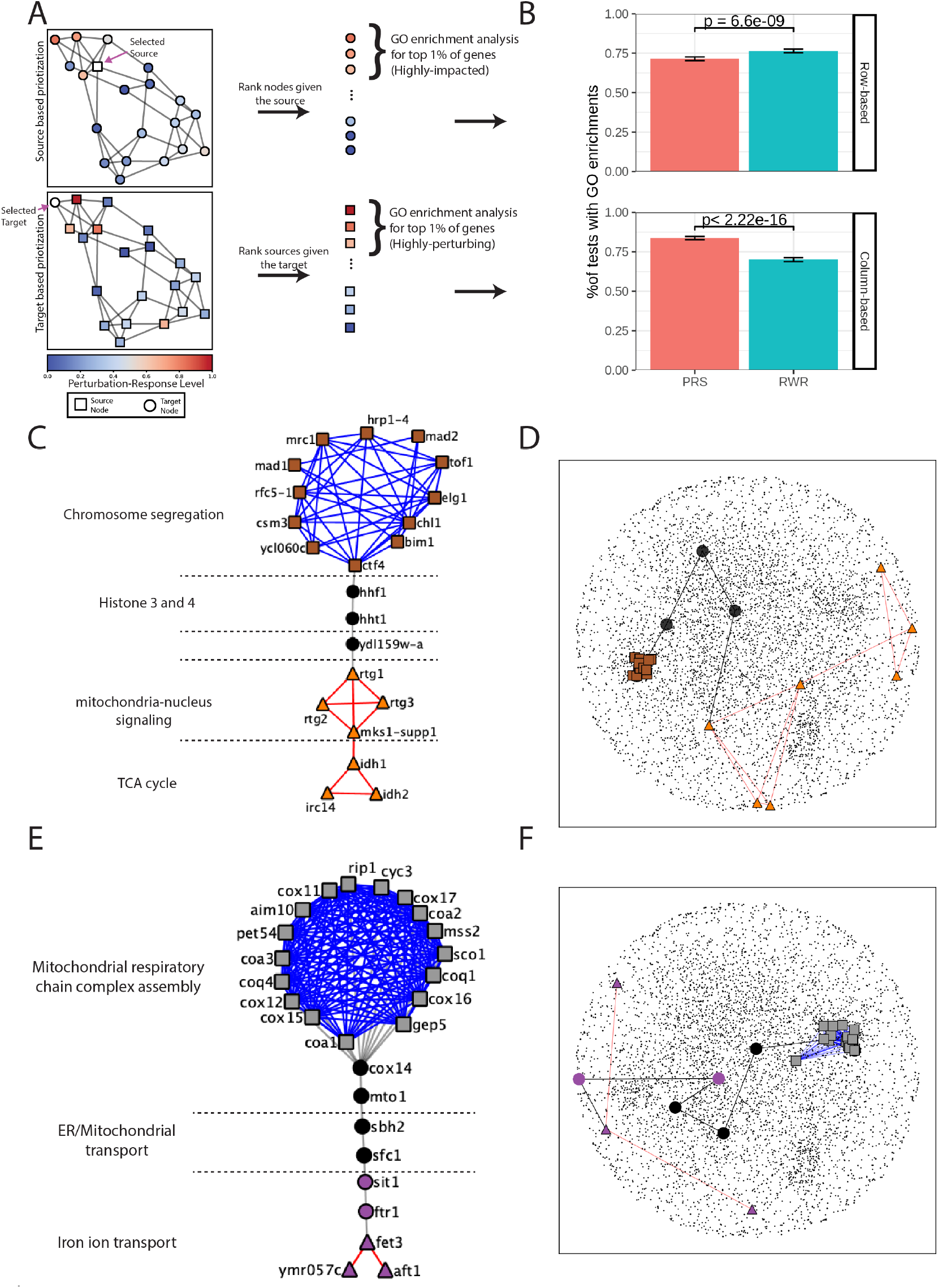
PRS identifies biologically meaningful long-range interactions without relying on prior knowledge. A) Graphical representation of the pipeline used to define highly responsive and highly influential gene sets. Each gene was analyzed as either the source of perturbation or responding to a perturbation and the genes were then ranked row-wise, representing the responding genes, and column-wise, representing the perturbed genes. Then for each gene, the top 1% of row-wise (responding) and column-wise (perturbing) genes were classified as highly responsive or highly influential set and GO enrichment analysis was performed. B) Comparison of the GO enriched ranked groups using PRS or random walk with restart (RWR) derived relationships for row-based (highly responsive sets) and column-based (highly influential sets) rankings, respectively. Error bars show 95% confidence intervals and *p*-values were calculated using two proportion Ztest) C) The PRS path from effector genes involved in chromosome segregation to sensors involved in mitochondria-nucleus signaling/TCA cycle which was found by perturbing the effector gene CTF4 and calculating the maximum information flow to sensor gene RTG1. D) Representation of the path shown in (C) within the GI PSN. E) The PRS path starting from effectors involved in respiratory complex assembly to iron transport sensors. Perturbation was applied to gene COA1 perturbation signal was followed through FET3. F) Representation of the path shown in (E) within the GI PSN.

Most of the ‘highly responsive’ and ‘highly influential’ gene sets were enriched in specific biological processes (72% and 84%, respectively; GO enrichment analysis, FDR<10%). Notably, most of the genes in ‘highly responsive’ or ‘highly influential’ sets had no direct interaction with the influential gene or the responding gene, demonstrating PRS’s ability to detect long-range relationships. Applying the same strategy to a similarity matrix derived from a random walk with restart (RWR) process, we found 77% and 70% of gene sets showed GO term enrichments for ‘highly responsive’ or ‘highly influential’ sets, respectively (Figure 5B, at FDR<10%). These results demonstrate that long-range interactions in the GI PSN harbor biological significance.

Most (70%) of the GO enriched ‘highly influential sets’ identified by our novel target based prioritization strategy contained at least one out of the 52 previously described effector genes. Interestingly, these effector-containing groups were mostly distinct with respect to the three effector clusters defined in Figure 4B: when a ‘highly influential set’ contained an effector from one of the three effector clusters, there were no effectors belonging to the other two clusters. These observations suggest that the three effector clusters influence different parts of the GI PSN.

As effectors and sensors are, by definition, critical nodes for long-range interactions, we inspected the information propagation paths derived from PRS signal transfer between clusters of effectors and sensors. A PRS path was defined as the node-weighted shortest path, where the weights were the inverse of the node responses to the perturbed node. This procedure identified the cellular pathways connecting effector and sensor clusters. For example, the effectors related to chromosome segregation and sensors related to the TCA cycle and mitochondria-nucleus signaling were found to be interconnected via histone modification genes (Figure 5C-D). Similarly, respiratory complex assembly effectors were connected to iron transport sensors via mitochondrial and ER transport genes (Figure 5E-F). Our analyses uncovered these otherwise buried paths as the long-range interactions between effector and sensor clusters, which are likely to constitute the pillars of the higher order organization of the GI PSN.

## Discussion

In this study, we adapted the PRS methodology, initially designed for characterizing allosteric signal transductions in molecular structures,^21–23^ to define the information propagation potential of genes in the yeast GI PSN. This approach identified clusters of critical effector and sensor genes representing different cellular processes and successfully detected long-range biological relationships between these distinct clusters. While effectors could have been estimated using other network centrality measures, such as degree, to our knowledge our approach is the only one able to sort the most critical effectors and pinpoint critical clusters of low-degree sensor genes.

Interestingly, the GI PSN demonstrated a superior propensity for information propagation compared to random networks with the same degree distribution. This suggests that other topological features of the GI PSN have evolved to enhance its capabilities for information sensing and transmitting, or overall functionality. Such features may include the hierarchical organization of increasingly more connected clusters previously described for GI PSN^28^ as well as the antenna-shaped motifs we discovered. These antenna-shaped motifs lack strong connections to the rest of the network, whereas effector clusters are tightly connected to the rest of the network. These patterns of assortative connections may reflect an evolutionary optimization of sensing properties for activating selected responses, while enhancing downstream cooperativity via effector genes.

Our results suggest that the precise topology of the GI PSN creates an opportunity or evolutionary adaptation for communication between distinct cellular processes. Beyond guilt-by-association^11^ and local network context analyses,^1^ our work illuminates how genes can communicate and affect processes beyond their local neighborhood. Altogether, our analyses add to the evidence^26,27^ that spring-based physical modeling of the networks can be a powerful tool to uncover the higher-order organization of the cell. It follows that more insight will arise from future work modeling biological networks as physical 3D objects. We anticipate that PRS strategies will extend to other types of complex networks, e.g., social, economic, microbiome where the identification of effectors and sensors together with the PRS paths may reveal important communication hubs and lines.

## Materials and Methods

### Yeast genetic interaction profile similarity network

We obtained the data from TheCellMap^32^ (https://thecellmap.org/costanzo2016/, file: Genetic interaction profile similarity matrices). Details of the network construction can be found in the supplementary materials of Costanzo et al.^28^ under the “Constructing genetic interaction profile similarity networks” section. In brief, the genetic interaction profile similarity between gene *i* and gene *j* is the Pearson’s correlation coefficient (PCC) between the genetic interaction profile vectors of *i* and *j*, which consist of genetic interaction scores experimentally estimated for all possible double mutants involving gene *i* or gene *j*:

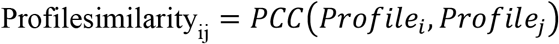

We used a PCC cutoff of 0.2 following the original publication,^28^ and derived the GI PSN containing every gene with at least one profile similarity of PCC > 0.2. This resulted in a network with 5,272 nodes and 39,866 unweighted and undirected edges.

### Elastic network models and perturbation response scanning matrix

We used the Gaussian network model (GNM) to represent the GI PSN as an elastic mass-and-spring network object. The overall connectivity of the network is represented by a Laplacian (also called Kirchhoff) matrix, whose diagonal elements are the degree of each node, and non-zero, negative off-diagonal elements (equal to −1) indicate the connected pairs of nodes. We first took the largest connected component of the GI PSN, which was represented by a GNM of *n* = 5,183 nodes and 39,816 edges. The corresponding Laplacian was used to perform the PRS analysis as described by Li et al.^33^ Mainly, we used *calcPerturbResponse* function in *ProDy*,^34^ a Python API designed originally for analyzing protein dynamics, to calculate the PRS matrix. This function first calculates the covariance matrix (Cov) between pairs of nodes, using the eigenvalues and eigenvectors of the Laplacian, followed by the normalization of each row upon dividing it by the diagonal element. The *ij*^th^ element of the resulting PRS matrix shows the response of the *j*^th^ node when the *i*^th^ node is perturbed. The row and column averages of the PRS matrix give the effectiveness and the sensitivity profiles as a function of gene index [1, *n*], respectively.

### PRS matrix clustering

To perform the clustering of the PRS matrix elements, we used a hierarchical clustering algorithm implemented in the Python package *SciPy*. We first capped the outliers in the PRS matrix by normalizing the values above 95% of the matrix to be equal to 95% value. Then we calculated the pairwise standardized Euclidean distance between genes using rows or columns of the PRS matrix as the coordinates, and used *ward* linkage metric to construct a dendrogram of the genes.

### Network properties

The following definitions are used. Node degree is the number of edges of a given node. Average neighbor degree is the average degree of the first neighboring nodes of a given node. Ratio of in-between edges for a given group of nodes is the ratio of the total number of edges that are directly connecting the nodes in the group to the total number of edges the nodes in the group have.

### Network rewiring

To rewire the network while keeping the degree distribution the same, we applied an edge swapping procedure. A swap between two randomly selected edges is accepted if the network connectivity is not violated, i.e., no network node is disconnected from the network, and if the newly generated edges are not already in the network. This process is repeated a minimum of 10 times the number of edges in the network. The resulting rewired network maintains the same degree for each node as the original network, but has different connections. For this process, we used *connected_double_edge_swap* function of the Python network analysis package, *networkx*.^35^

### Gene ontology enrichment analyses

GO trees and annotations were downloaded from http://geneontology.org/ on May 20, 2021. We used the Python package, *GOATools*,^36^ to calculate the number of genes associated with each GO term in the study group and the overall population of (all) genes. We excluded the evidence codes ND (no biological data available), IGI (inferred from genetic interaction), and HGI (inferred from high throughput genetic interaction) to remove any associations originating from the genetic interaction network we used. We applied Fisher’s exact test and false discovery rate (FDR) multiple testing correction to calculate corrected *p*-values for the enrichment of GO term in the study group. FDR<0.1 was taken as requirement for significance.

### Sensors and effectors group comparisons

Kruskal-Wallis test was used to statistically investigate the differences between effector or sensor groups in terms of their degree, effectiveness or sensitivity values for the analyses shown in Figure 3D-E and Figure 4C-D. We applied *kruskal.test* function in R with a significance level of *α* = .05. To find the group that deviates from the null model, we used Tukey’s HSD test,^37^ which is equivalent to a pairwise Wilcoxon test with multiple testing corrections.

### Random walk with restart

We used the RWR formula defined in Leiserson et al.^16^ We calculated steady-state solution of RWR for each node. Then, we created an RWR matrix where each row *i* represents the steady-state solution vector for RWR starting at node *i*. While this is similar to the PRS, row sums of the RWR matrix equal to one, showing the probability distribution of each random walk process and column sums are the PageRank centrality. RWR thus could not have been used instead of PRS to identify effectors and sensors in the network.

### PRS (or RWR) ranking

To define highly responsive and highly influential gene sets, we implemented the ranking strategy illustrated in Figure 5A. For each gene *i*, we took the *i*^th^ row of the PRS (or the RWR) matrix, sorted it in descending order, and took the top 52 genes as highly responsive group. Similarly, for each gene *j*, we took *j*^th^ column of the PRS (or the RWR) matrix then ranked and selected in the same way to define highly influential group. Then we used *GOATools* to calculate the enriched GO terms corresponding to these groups of 52 genes as explained above.

### PRS path analysis

For each path starting at gene *i*, we took *i*^th^ row values of the PRS matrix as node weights. To find the path that carries the maximum information, we inversed node weights and used Dijkstra’s algorithm to find the shortest weighted path. Cytoscape and *networkx* were used to visualize the paths between effectors and sensors. Annotations were done manually using gene descriptions in *Saccharomyces* Genome Database (SGD).^38^

## Acknowledgments

The authors are grateful to April Rich, Dr. Branden Van Oss, Dr. Aaron Wacholder, Carly Houghton, Jiwon Lee, Lin Chou, Dr. Saurin Bipin Parikh for reviewing the manuscript prior to submission.

## Conceptualization

O.A., S.Z., I.B., and A.-R.C.; Methodology: O.A., S.Z.; Investigation: O.A.; Writing-original draft: O.A.; Writing-review and editing: O.A., S.Z., I.B., and A.-R.C.; Supervision: I.B. and A.-R.C.

## Source files and code

All source code and csv files for figure generation are accessible online at https://www.github.com/oacar/enm_package

